# Polarity sorting of actin filaments by motor-driven cargo transport

**DOI:** 10.1101/2024.06.24.600553

**Authors:** Oghosa H. Akenuwa, Steven M. Abel

## Abstract

During the active transport of cellular cargo, forces generated by cargo-associated molecular motors propel the cargo along cytoskeletal tracks. However, the forces impact not only the cargo, but also the underlying cytoskeletal filaments. To better understand the interplay between cargo transport and the organization of cytoskeletal filaments, we employ coarse-grained computer simulations to study actin filaments interacting with cargo-anchored myosin motors in a confined domain. We show that cargo transport can lead to the segregation of filaments into domains of preferred filament polarity separated by clusters of aggregated cargoes. The formation of polarity-sorted filament domains is enhanced by larger numbers of cargoes, more motors per cargo, and longer filaments. Analysis of individual trajectories reveals dynamic and heterogeneous behavior, including locally stable aggregates of cargoes that undergo rapid coalescence into larger clusters when sufficiently close. Our results provide insight into the impact of motor-driven organelle transport on actin filaments, which is relevant both in cells and in synthetic environments.

**SIGNIFICANCE**
The actin cytoskeleton is vital for intracellular transport, and there is an intricate interplay between the organization of the actin network and cargo transport by molecular motors. In this work, we use computer simulations to demonstrate that the transport of cargoes by teams of molecular motors can lead to the emergence of filament domains, separated by clusters of cargoes, that are sorted by the polarity of filaments. The results, which highlight feedback between transport and filament organization, provide insight into mechanisms influencing the cytoskeleton in cells and in reconstituted systems.

## INTRODUCTION

Active, motor-driven transport is essential for proper cell function and growth in eukaryotic cells (1). It is carried out by molecular motors moving in preferred directions along cytoskeletal filaments, each of which has a defined polarity (1, 2). For cellular cargoes such as organelles and vesicles, transport is often carried out by multiple cargo-associated motors working in tandem (3). Cytoskeletal filaments serve as tracks for molecular motors, yet active transport of organelles also impacts the organization of the cytoskeletal network (4–7). Thus, there is feedback between motor-driven cargo transport and the underlying organization of the cytoskeletal network. However, much remains unknown about how organelle transport impacts cytoskeletal organization. Better understanding the interplay between transport and network organization would yield insights into regulation of cellular transport and design principles for reconstituted and synthetic systems.

Systems of molecular motors and cytoskeletal filaments have been studied *in vivo, in vitro*, and *in silico* (8–11). The most extensive studies have been reconstituted systems and associated computational studies, which have focused primarily on systems of myosin II motors with actin filaments or systems of kinesin motors with microtubules, typically in reconstituted quasi-2D environments (12–16). Cytoskeletal structures observed in these systems include dense asters, nematic lanes, vortices, and flows (15, 17). Forces generated by molecular motors can cause active sliding of filaments and lead to local sorting, with filaments locally aligned in a polar manner (18, 19). Sorting by filament polarity has been associated with polar flows, vortices, and asters in kinesin-microtubule systems (18, 19) and asters in myosin II-actin systems (20, 21). Polarity sorting in these systems has been attributed to the end-dwelling nature of kinesins and myosin II (20, 22). Recently, Memarian et al. (23) studied microtubules propelled by membrane-bound kinesin motors. The motors, which organized the microtubules into locally-aligned nematic lanes, were concomitantly reorganized because of their lateral mobility.

Computational approaches have provided important insights into the physical mechanisms underlying the organization and dynamics of motor-filament systems. A successful class of models uses dynamical, coarse-grained representations of filaments and motors, with filaments modeled as semiflexible polymers and motors modeled either explicitly with heads that bind to and move along filaments (24, 25) or implicitly as tangential forces acting along the contour of filaments (26, 27). Simulation packages such as CytoSim (28), MEDYAN (29), AFINES (30), and aLENS (22) have been developed and utilized to study systems of myosin II motors and actin filaments. Simulations using these approaches have elucidated collective behavior of filaments in response to motor forces, including the formation of bundles (31, 32), asters (24, 31), vortices, and rings (27, 33). Computational work focused on the transport of organelles has focused largely on the impact of multiple motors transporting an organelle along a single cytoskeletal filament. These studies have characterized how properties such as the number of motors bound to the organelle impact its motion along a filament (34–38).

While we are interested in general insight into the interplay between cargo transport and filament organization, we are motivated specifically by understanding transport and cytoskeletal organization in plant cells. In plants, there are two main classes of motors involved in organelle transport: myosin VIII and myosin XI. Myosin XI has also been shown to be involved in the organization of the actin cytoskeleton. In pollen tubes, myosin XI knockout mutations lead to disruption of the actin network, reduced ordering of actin filaments in bundles, and reduced dynamic rearrangements of the actin network (5, 6, 39), while in root epidermal cells, myosin XI mutants exhibit increased bundling and ordering of actin filaments (7). Confinement also impacts the organization of actin networks (40, 41). In plant cells, the cell wall provides rigid external confinement, and the large vacuole can occupy up to 90% of the cytoplasm, leading to quasi-2D environments between the vacuole and plasma membrane (42). Recent experimental and computational studies have demonstrated that confinement impacts the organization of motor-filament systems, promoting the formation of polar and nematic flows (43–45), as well as aster-like structures (22).

Despite a robust body of prior work, the interplay between motor-driven organelle transport, spatial confinement, and cytoskeletal organization remains largely unexplored. Previously, we investigated the impact of crosslinking proteins on actin organization in confined geometries, focusing only on crosslinking proteins and filaments to obtain insight into effects of crosslinkers (7, 41). In this work, we study the impact of cargo transport by molecular motors on the organization and dynamics of actin filaments in confined environments. We take a similar approach and focus on systems consisting only of cargo-associated molecular motors and filaments, which allows us to gain insight into the effects of motor-driven cargo transport without other confounding effects. This reveals physical mechanisms relevant in cellular settings and may also provide direct insight into reconstituted experimental systems.

## METHODS

We used the Actin Filament Network Simulation (AFINES) model, developed by Freedman et al. (30, 46, 47), which is a coarse-grained model that uses kinetic Monte Carlo and Brownian dynamics methods to simulate actin filaments and molecular motors. We provide a brief overview of the method, details of which can be found in Ref. (30), and discuss how we implemented cargo transport. A schematic representation of the model is shown in Figure 1.

**Figure 1:**
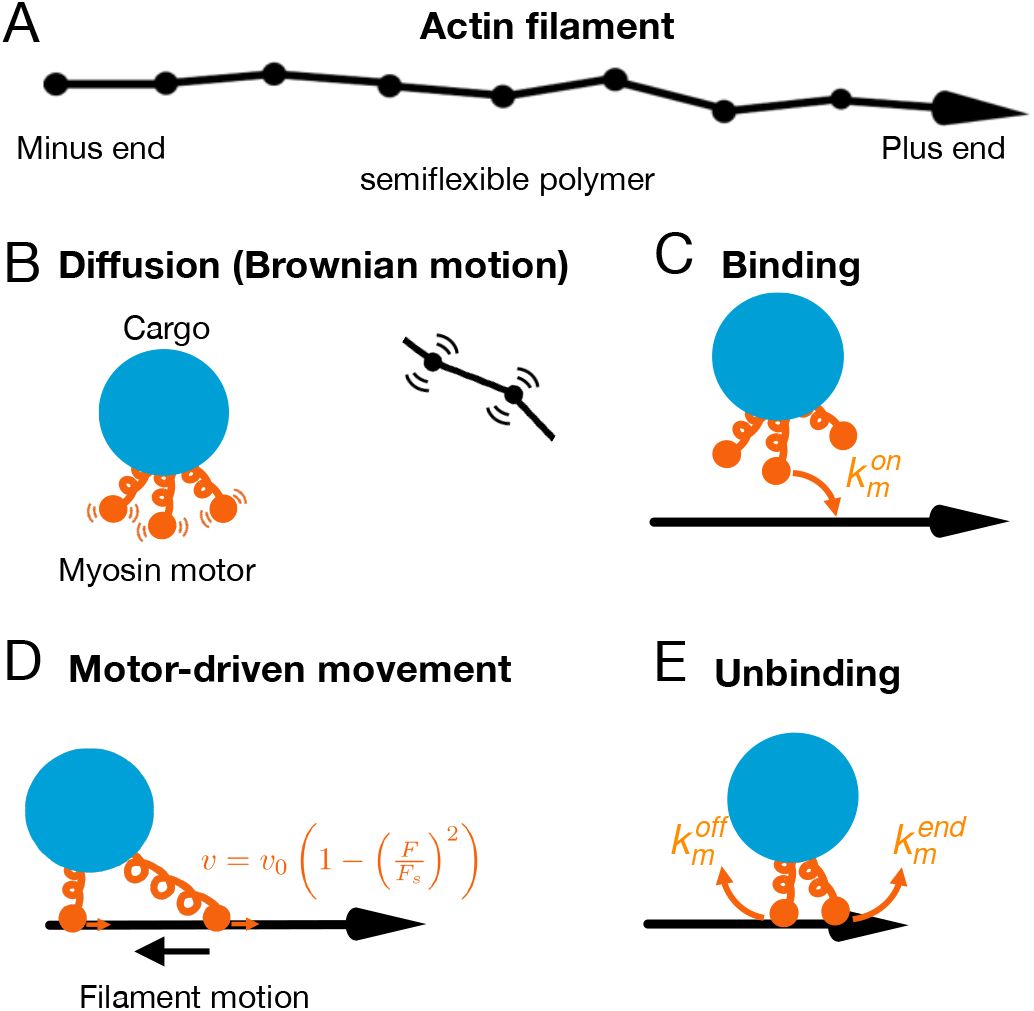
Schematic describing the cargo-motor complex and actin filaments. A cargo is shown in blue and motors in orange. Motors are attached to a common point at the center of the cargo.

In the AFINES model, actin filaments are modeled as semiflexible, bead-spring polymers in two dimensions. One end of the filament represents the plus (barbed) end and the other represents the minus (pointed) end (Fig. 1A). Excluded volume interactions are neglected. The bending modulus is chosen so that the persistence length of an isolated actin filament is 17 μm. Cargoes are modeled as a spherical beads of radius *r*_*c*_ = 250 nm which undergo Brownian motion with a drag constant *η* = 6*πvr*_*c*_, where *v* is the dynamic viscosity of the medium (Fig. 1B). Motors are treated as springs with one end permanently attached to a cargo and the other end allowed to bind and unbind filaments (Fig. 1C). The potential energy of a motor is given by 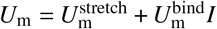, where *I* is 1 when the motor is bound to a filament and 0 otherwise, and

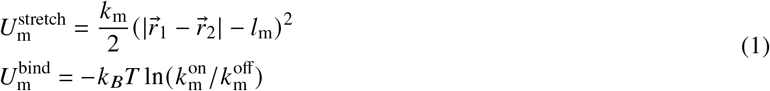

Here *l*_m_ = 350 nm is the equilibrium length of the motor, *k*_m_ = 10 pN/μm is its stretching stiffness, 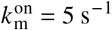 is the binding rate, and 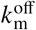 is the unbinding rate. We assume that the motors are attached to a common point at the center of the cargo, and thus *l*_m_ represents the distance from the center of the cargo to the filament-binding end of the motor.

When a motor is bound to a filament, it travels toward the plus end with a load-dependent velocity (Fig. 1D) given by

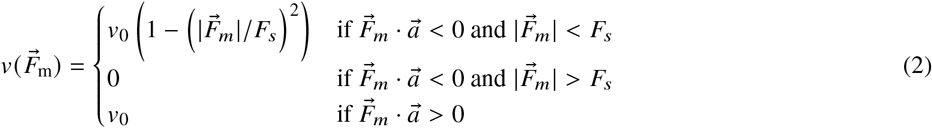

where *v*_0_ = 7 μm/s is the unloaded velocity, 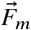 is the tensile force on the motor, *F*_*s*_ = 0.5 pN is the stall force of the motor, and 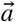 is a displacement vector along the filament link to which the motor is bound pointing toward the plus end. It has been shown that forward loads on motors like myosin V do not significantly impact their velocity, but backward loads slow them down (48, 49). Hence, we assume that the velocity is impacted only when the force is directed away from the direction of travel. Motors unbind from filaments with a rate 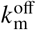 (Fig. 1E). We assume that 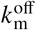 is load sensitive and given by

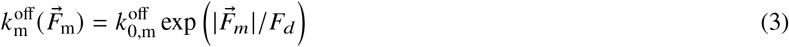

where 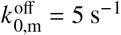 is the off rate of the motor in the absence of force and *F*_*d*_ = 1 pN is the characteristic detachment force of the motor. This slip-bond form of the off-rate has been demonstrated experimentally for kinesin motors (50), and we assume similar behavior for the motors here.

Tensile forces from motors are propagated to the center of mass of the associated cargo and, using the lever rule, to filament beads neighboring a bound filament segment. We assume that motors rapidly dissociate upon reaching the plus end (51) and set the unbinding rate of motors at the plus end to 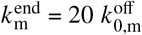 (Fig. 1E). Excluded volume of motors and cargoes is ignored.

Dynamics are governed by a kinetic Monte Carlo algorithm to update the binding and unbinding of the motors and Brownian dynamics to update the positions of the filaments, motors, and cargoes. In a given time interval Δ*t*, each unbound motor binds with probability 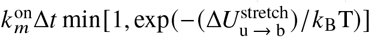, and each bound motor unbinds with probability 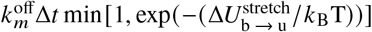. Here 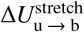 represents the difference in stretching energy of the motor between the unbound and bound state. Similarly, 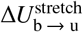 represents the difference in stretching energy between the bound to unbound state. Then, given the updated state of the system, positions of the filament beads, cargoes, and motors are updated using overdamped Langevin dynamics, propagating the time forward by Δ*t*. Bound motors move a distance *v*Δ*t* along the filaments towards the plus end. The process is then repeated.

### Parameters studied and initialization

We considered filament lengths of 10 and 5 μm. The number of filaments (*N*_*f*_ = 100, 200) was chosen to keep the total filament length constant across all simulations (1000 μm). The number of cargoes was varied, with *N*_*c*_ = 10, 20, 40, and 80. The number of motors per cargo was varied, with *N*_*m*_ = 1, 2, 4, and 8. The system shape was a rectangular box of size 40 μm × 10 μm. These dimensions are typical of quasi-two-dimensional environments found in plant cells and are accessible in reconstituted systems (52–54). Reflective wall boundary conditions were imposed. Table S1 provides a list of the parameters used in this study.

For each set of parameters, 50 independent simulation trajectories were generated and analyzed. Simulations were initialized by randomly distributing filaments in the simulation box and equilibrating the filaments without cargoes or motors. For the shorter filaments, we used a 500 s simulation run for equilibration. The longer filaments were slower to equilibrate, so we implemented a step-wise annealing procedure to facilitate the equilibration. The simulation consisted of the following temperatures and simulation times: 7240 K for 300 s, followed by 5790 K, 4350 K, and 3620 K for 150 s each, followed by 1810 K, 724 K, 543 K, 260 K, and 298 K (target temperature) for 300 s each. Unbound cargo-motor complexes were then added uniformly at random, and the simulations were run for 500 s.

## RESULTS

### Cargo transport leads to spontaneous organization

Figure 2 shows time-lapse snapshots of sample trajectories with 10 μm filaments, 20 cargoes, and varying numbers of motors per cargo (*N*_*m*_). Filaments are colored according to their initial orientation, which we characterize by the end-to-end vector connecting the minus to plus end. At *t* = 0 s, randomly distributed cargoes are added to a system of equilibrated filaments. Motors rapidly associate with filaments, leading to transport of cargoes and rearrangement of filaments. Notably, for sufficiently large *N*_*m*_, filaments organize over time, yielding regions enriched in filaments of a similar initial orientation. Here right-pointing filaments aggregate towards the left of the simulation box and left-pointing filaments aggregate towards the right. We refer to this as polarity sorting. In both domains, the plus ends tend to accumulate near the center of the system. Concurrent with the organization of filaments, the cargoes tend to cluster between the regions of opposite polarity. Thus, if a filament diffuses into the central region, motors will likely bind to it and generate a force pushing the filament away from the middle. Similarly, if a cargo diffuses away from the central region, its motors will likely engage filaments, leading to a net force directing the cargo toward the middle.

**Figure 2:**
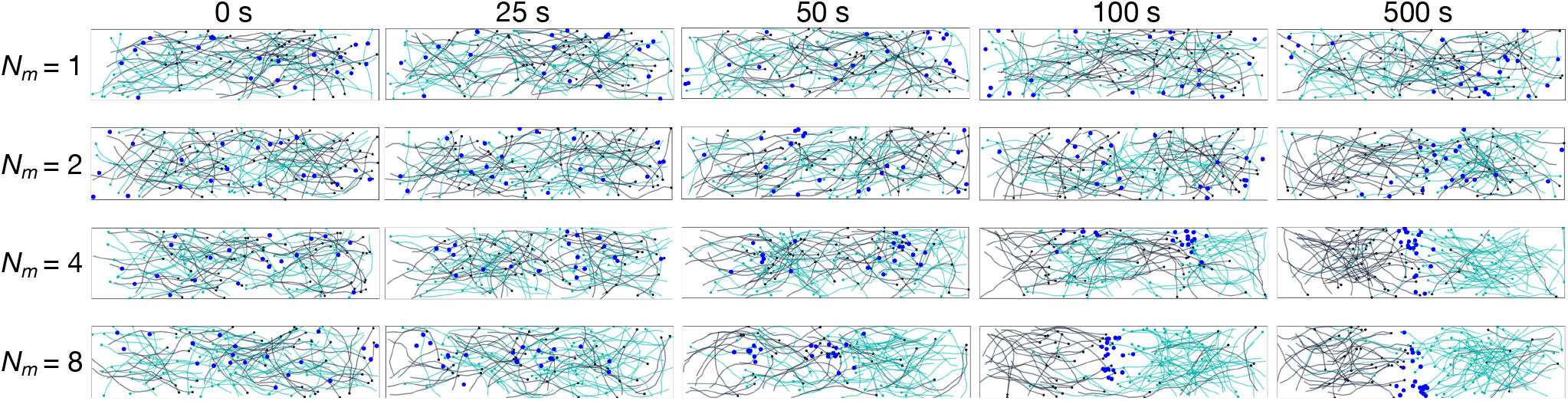
Time-lapse snapshots of trajectories with 10 μm filaments, 20 cargoes, and varying numbers of motors per cargo (*N*_*m*_). Filaments are colored according to their initial orientation relative to the long axis, as determined by the end-to-end vector connecting the minus to plus end. Left-facing filaments are shown in green and right-facing filaments in charcoal. Cargoes are depicted by blue circles. Individual motors are not shown. The system size is 40 μm × 10 μm.

### Transport by teams of motors promotes formation of domains

The individual trajectories in Figure 2 suggest that motor-driven cargo transport can lead to the spontaneous organization of filaments and cargoes. To further investigate, we characterized the spatial organization of filaments upon varying the number of cargoes (*N*_*c*_) and the number of motors per cargo (*N*_*m*_). We determined the mean filament occupancy as a function of position by dividing the simulation box into square voxels with a side length of 0.2 μm and determining the number of filaments in each voxel. We averaged this quantity over the last 50 s of 50 independent trajectories. We further characterized the mean filament occupancy as a function of the position (*x*) along the long dimension of the simulation box to facilitate a quantitative comparison.

Figures 3A and B show the distribution of filaments within the simulation domain. When each cargo has only a single motor (*N*_*m*_ = 1), the distribution of filaments is characterized by a single, extended region of high occupancy. The filaments appear homogeneously distributed in the central part of the simulation domain, and the density is depleted near the edges of the domain. The 1D filament distributions show a uniform distribution in the central part of the long axis with steady decline towards the edges; the distribution is similar to the case with no motors. Depletion near the edges is a feature of all systems considered because there is an effective repulsion of filaments from the walls (55, 56).

**Figure 3:**
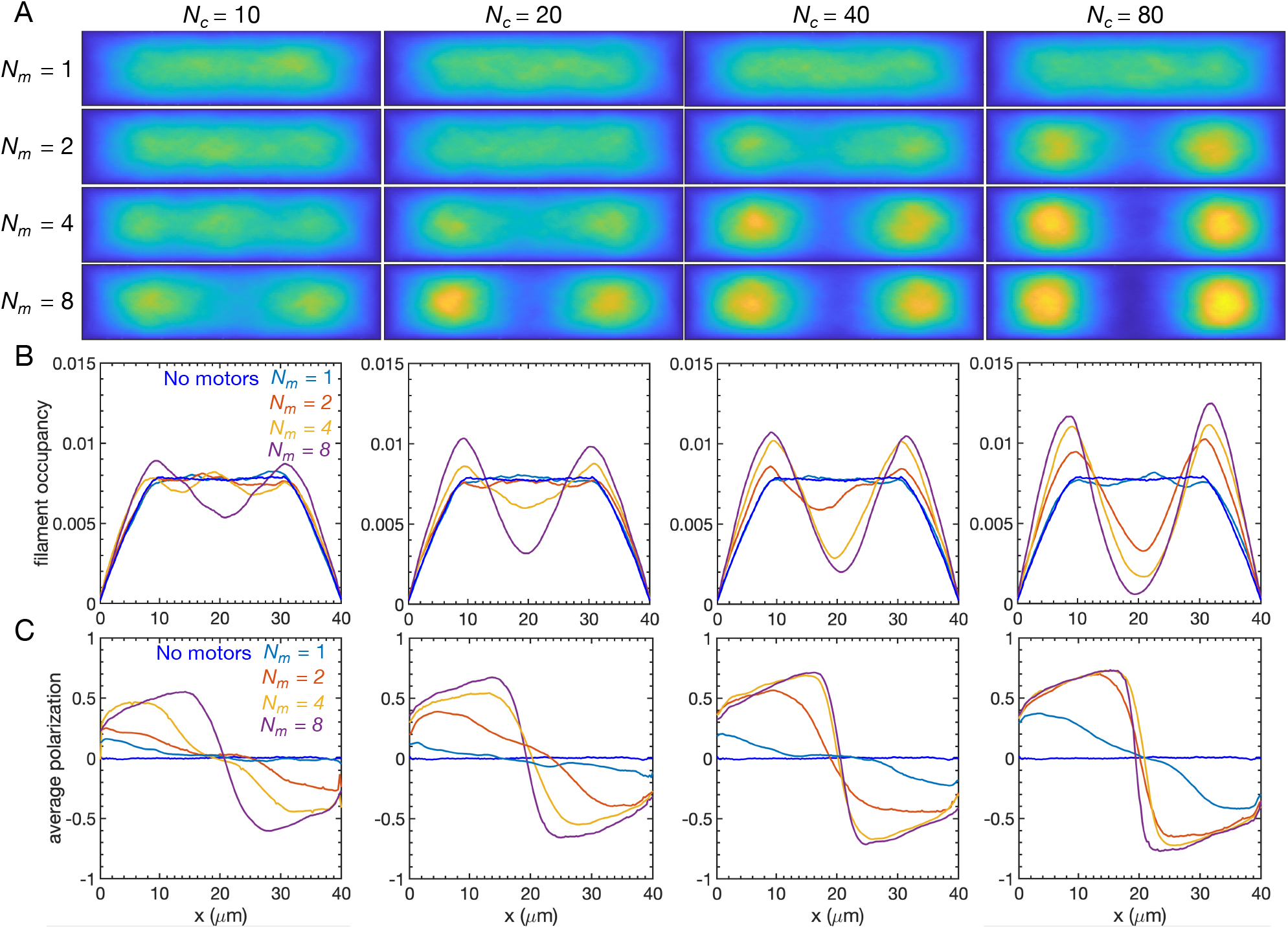
(A) Spatial distribution of filaments within the simulation domain (40 μm × 10 μm). The mean filament occupancy is shown for 10 μm filaments and different values of *N*_*c*_ and *N*_*m*_. All heatmaps use the same color scale and can be compared directly; yellow denotes high occupancy. (B) Corresponding 1D distribution of filaments along the long dimension of the simulation domain (*x*). Each column corresponds to the value of *N*_*c*_ shown in (A). (C) Average polarization in the *x*-direction. Each heatmap and associated curve is an average over the final 50 s of 50 trajectories.

When more than one motor is associated with each cargo, multiple filament domains emerge, as demonstrated by regions of high filament occupancy separated by a region of low occupancy. Spatially segregated domains are promoted by increasing *N*_*c*_ and/or *N*_*m*_. On average, filaments tend to accumulate in regions that are off-center of the long axis of the system. As *N*_*c*_ and *N*_*m*_ increase, the regions become more pronounced, and the separation between the regions increases. Note that it is not simply the total number of motors that governs the filament distribution. For example, for the cases with 80 total motors (plots along the anti-diagonal in Figure 3A), fewer cargoes with more motors per cargo result in more pronounced and separated filament domains.

To characterize the polarity of the filaments, we determined the average polarization in the *x*-direction as a function of *x*(Fig. 3C). For each region of thickness Δ*m* = 0.2 μm, we determined the unit vector pointing from the minus to plus end of each 0.1 μm segment of filament in the region and averaged over all such vectors. Positive values indicate regions in which plus ends of filaments point to the right on average; negative values indicate regions in which they point to the left on average. Each distribution is an average over the last 50 s of 50 trajectories.

For the case with no motors, the average polarization is zero everywhere, indicating no net bias in the filament polarity in the absence of motors. As *N*_*c*_ and *N*_*m*_ increase, the average polarization illuminates two distinct regions. The region left of center has positive values, indicating filaments with their plus ends pointing to the right. The region right of center has negative values, indicating filaments with their plus ends pointed to the left. This confirms that the filament domains shown in Figs. 3A and B correspond to regions of filaments with distinct polarity. Furthermore, the magnitude of the polarization in the two regions increases as *N*_*c*_ and *N*_*m*_ increase, indicating enhanced sorting of the filaments. The decrease in the magnitude near the edges is likely a consequence of filaments that were not sorted before the cargo domain formed and the reorientation of filaments due to thermal fluctuations.

The emergence of filament domains coincides with a change in the spatial organization of cargoes (Figures 4A and B). With small numbers of cargoes and one motor per cargo, the cargoes remain uniformly distributed throughout the simulation domain. As *N*_*c*_ and/or *N*_*m*_ increase, the cargoes accumulate near the center of the long dimension. The region of increased residence coincides with the central region depleted of filaments.

**Figure 4:**
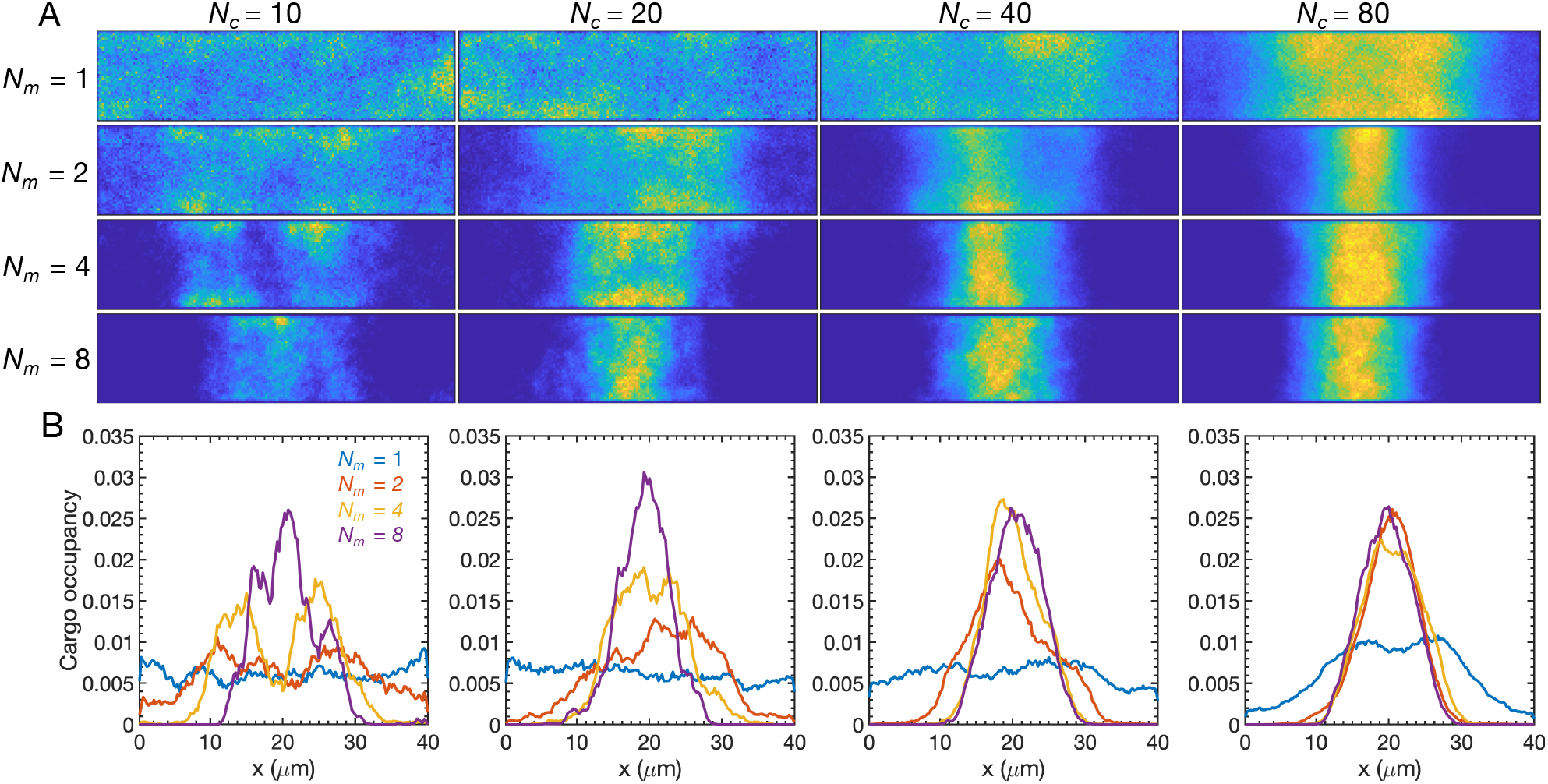
(A) Mean cargo occupancy within the simulation domain with 10 μm filaments. (B) Corresponding 1D cargo distribution along the long dimension of the simulation box (*m*).

The results above provide average measures of organization, but stochasticity leads to differences between trajectories. Figure 5 shows the final snapshots of sample trajectories with *N*_*m*_ = 8 and different numbers of cargoes. The snapshots demonstrate polarity sorting, but the location of the clustered cargoes varies between trajectories. To further characterize differences between trajectories, Figures S1 and S2 show the filament and cargo occupancy distributions for each independent trajectory. There is marked heterogeneity in the distributions, but individual trajectories more closely resemble the average distribution for large values of *N*_*c*_ and *N*_*m*_. In these cases, a large cluster of cargoes remains near the center of the domain with strong filament sorting on each side.

**Figure 5:**
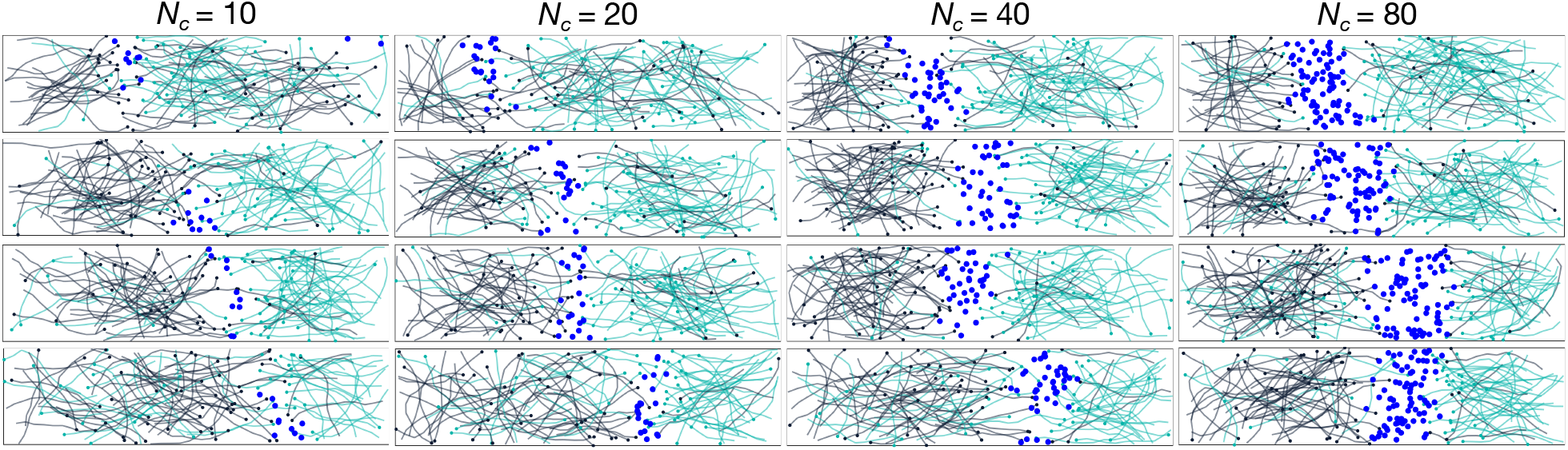
Final snapshots of independent trajectories with 10 μm filaments and *N*_*m*_ = 8. Four representative replicates are shown for each value of *N*_*c*_. Filaments are colored according to their initial orientation, with left-facing filaments shown in green and right-facing filaments shown in charcoal. Cargoes are depicted by blue circles. Individual motors are not shown.

### Dynamics of polarity sorting and cargo aggregation

To investigate the dynamics of domain formation and polarity sorting, we characterized the net displacement of filaments of similar initial polarity. Specifically, we calculated the relative change in the mean center of mass of right-pointing filaments as a function of time (Figure 6A). For *N*_*c*_ = 10 and *N*_*m*_ = 1, there is no pronounced change in the average filament location. Increasing *N*_*c*_ and/or *N*_*m*_ leads to a larger change in the center of mass, indicating net displacement of the filaments toward the left. Left-facing filaments exhibit similar net displacement to the right. The net displacement demonstrates that polarity sorting is faster and more pronounced for larger values of *N*_*c*_ and *N*_*m*_. For smaller values of *N*_*c*_ and *N*_*m*_, the net displacements have not plateaued, indicating a slow, ongoing process of filament transport by the motor-driven motion of cargoes. There is a slight increase in the value of the net displacement at long times for cases that plateau at early times; this is associated with small numbers of filaments changing their orientation and being transported across the cargo cluster.

**Figure 6:**
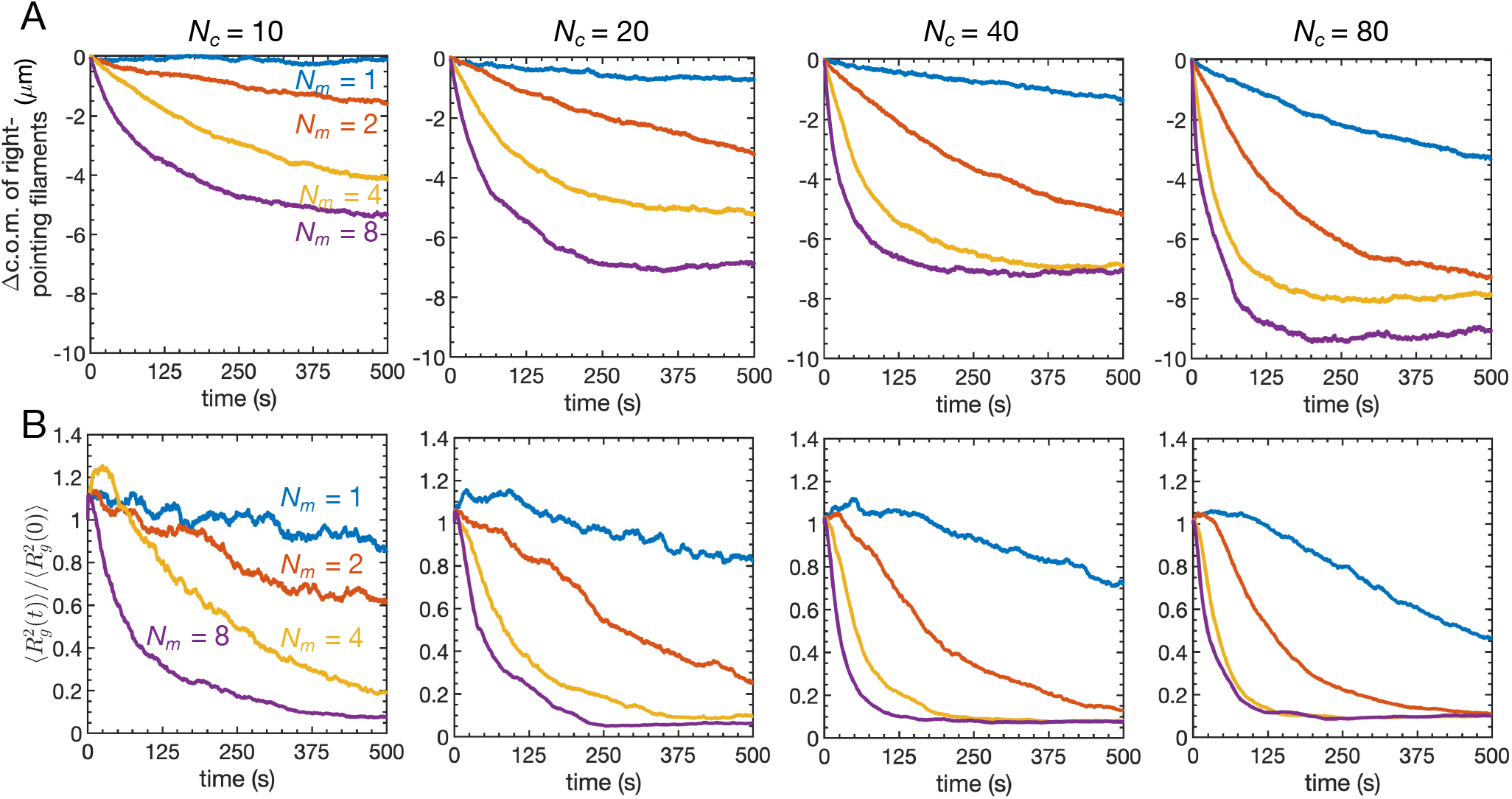
(A) Change in the center of mass (c.o.m.) of right-pointing filaments. (B) Mean square radius of gyration of cargoes normalized by the initial mean square radius of gyration. Each curve is an average over 50 trajectories.

We further characterized the dynamics of cargo aggregation by determining the radius of gyration (*m*) of the centers of mass of the cargoes. Because the cargoes are initially distributed uniformly at random, the average radius of gyration decreases when they aggregate into a cluster (Figure 6B). Increasing *N*_*c*_ and/or *N*_*m*_ leads to a faster, more pronounced decrease in the normalized radius of gyration, indicating that more cargoes and motors promote aggregation. With one or two motors per cargo, the radius of gyration continues to decrease at 500 s, indicating a slow and ongoing reorganization of the system induced by transport of cargoes.

Individual trajectories provide additional insight into the dynamics of cargo aggregation (see Movie S1 for an example with *N*_*c*_ = 40 and *N*_*m*_ = 8). Figure 7A shows the normalized radius of gyration as a function of time for individual trajectories with *N*_*c*_ = 80 and *N*_*m*_ = 8. Additional cases are shown in Figure S3. The trajectories highlight two dynamical scenarios. In the first, the radius of gyration decreases rapidly to a steady state value consistent with a single cluster. In the other scenario, the radius of gyration first rapidly decreases to a plateau-like or slowly decreasing intermediate regime. This is followed by a sharp and rapid decline to the steady state value consistent with a single cluster.

**Figure 7:**
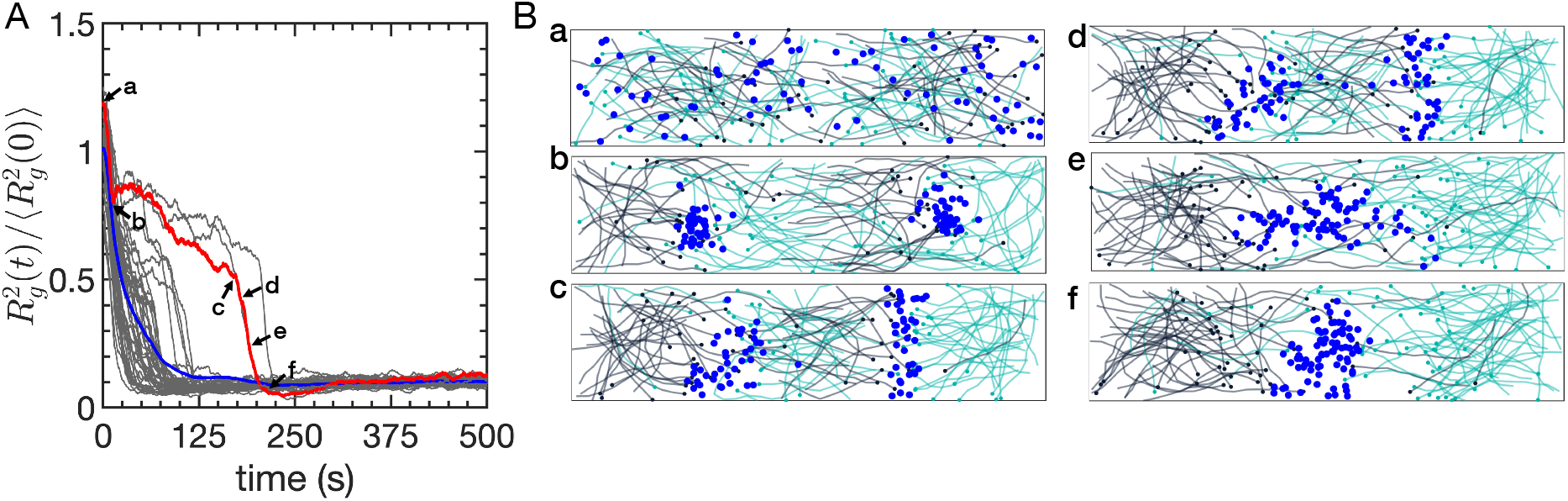
(A) Normalized square radius of gyration of cargoes. Individual trajectories are shown as grey curves, with the mean value in blue. (B) Snapshots taken at labeled timepoints of the trajectory shown in red.

Figure 7B shows snapshots taken from a sample trajectory exhibiting dynamics consistent with the second scenario. The cargoes and filaments, which are initially dispersed, quickly aggregate into two clusters (snapshots a, b). The filaments around each cluster are locally sorted by polarity, causing the configuration to be locally stable. The location of the clusters drift closer to one another in the intermediate regime, leading to filaments of opposite polarity coexisting in the space between them (snapshot c). This is followed by rapid coalescence into a single cluster (snapshots d - f). The coalescence is driven by cargoes that are transported on filaments pointing away from the cluster of origin in the region between the two clusters. This transports the cargo toward the other cluster while also pulling the filament towards the original cluster, facilitating more cargoes being transported away from the original cluster and depletion of filaments in the region between the clusters. This results in aggregation of the cargoes into a single cluster, which persists over time. Figure S3 indicates that there is a population of trajectories for all cases with *N*_*m*_ *m* 2 that reach a value of *m*^2^ that is consistent with a single cluster of cargoes. Increasing *N*_*m*_ and *N*_*c*_ increases the size of the population and decreases the typical times for trajectories to reach the single-cluster state.

### Shorter filaments exhibit less pronounced domain formation

Because 10 μm filaments are similar in length to the size of the short dimension of the simulation domain, the filaments tend to partially align with the long dimension (41). In contrast, shorter filaments are impacted less significantly by the confinement. Additionally, shorter filaments are likely to generate ordering on shorter length scales and change their orientation on shorter time scales. To investigate the impact of filament length, we considered shorter (5 μm) filaments while increasing the number of filaments to keep the total filament length constant.

Figure 8A shows the average, spatially-resolved filament occupancy within the simulation box. For most values of *N*_*c*_ and *N*_*m*_, the filaments on average reside in a single domain that is homogeneous except for depletion of filament density near the edges. The distribution is similar to the case without motors (Fig. S4). When *N*_*m*_ and *N*_*c*_ are sufficiently large, regions of enriched filament density emerge near the ends of the long dimension. However, the high-occupancy regions are less pronounced and farther apart than the analogous case with 10 μm filaments (Figs. 3A and B). Thus, for shorter filaments, more cargoes and motors per cargo are required to observe significant aggregation of the filaments on average.

**Figure 8:**
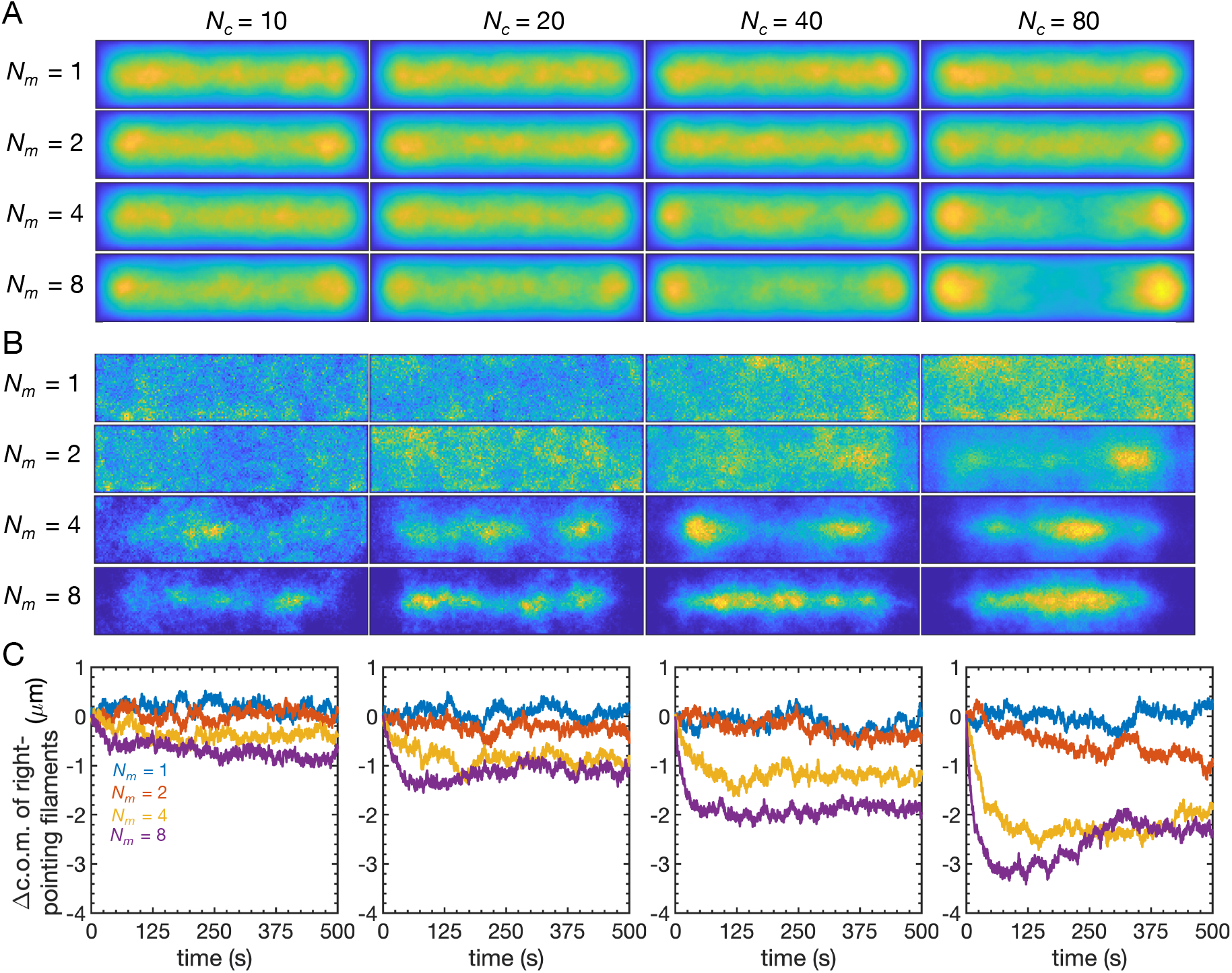
(A) Spatial distribution of filaments within the simulation domain for shorter (5 μm) filaments. The mean filament occupancy is shown for different values of *N*_*c*_ and *N*_*m*_. All heatmaps use the same color scale and can be compared directly; yellow denotes high occupancy. (B) Mean cargo occupancy within the simulation domain. (C) Change in the center of mass (c.o.m.) of right-pointing filaments.

Figure 8B shows the spatial distribution of cargoes. For sufficiently small valuers of *N*_*c*_ and *N*_*m*_, the cargoes remain uniformly distributed. As *N*_*c*_ and *N*_*m*_ increase, there is a depletion of cargo occupancy near the edges of the simulation domain and an enhancement near the center. In contrast with the longer filaments, the depletion of cargo density occurs in both dimensions, and not just along the longer dimension. This is because filaments are short enough to be ordered by polarity sorting in all directions. Figure 8C demonstrates that the spatial organization of filaments and cargoes is associated with the sorting of filaments by polarity. Increasing *N*_*c*_ and *N*_*m*_ leads to a larger change in the center of mass of right-pointing filaments, but the decrease is less pronounced than with 10 μm filaments.

Snapshots of individual trajectories show localized clusters of cargoes, which become more pronounced as *N*_*c*_ increases (Figure S5). It is notable that in cases with sufficiently large *N*_*m*_ and *N*_*c*_, individual trajectories exhibit more localized cargo clusters (Figs. S6 and S7) than is evident in the average distribution (Figure 8B). The relatively uniform average distribution results from the broad range of locations of individual clusters across different trajectories, and the fact that cargo clusters move over time. Further, for *N*_*m*_ = 4 and 8, a slow decrease in the average radius of gyration at long times appears to be driven in part by the rare coalescence of cargoes into a single, stable cluster. The emergence of such clusters occurs in a narrower range of parameter space and is less common than with longer filaments, indicating that larger values of *N*_*c*_ and *N*_*m*_ are needed to generate a single stable cluster.

## DISCUSSION

Although cytoskeletal filaments serve as tracks for motor-driven transport, forces generated by molecular motors can also impact the organization of the cytoskeleton, leading to feedback between transport and organization. In this work, we used computer simulations to study the impact of motor-driven cargo transport on the organization of actin filaments in confined domains. Figure 3 illustrates our key results and demonstrates the emergence of distinct filament domains separated by a domain enriched in cargoes. The filament domains are characterized by distinct polarity, with the plus ends of filaments pointing toward the cargo domain. By varying parameters in the simulations, we demonstrated that polarity sorting is promoted by increased numbers of cargoes, more motors per cargo, and longer filaments.

Physically, as motors walk toward the plus end of a filament, drag on the cargo serves as a resistive load and generates a tensile force on the motors. This force leads to motion of the cargo along the direction of the filament, but it simultaneously pulls the filament toward the cargo. Thus, even cargoes with a single motor can impact the organization of actin filaments. However, when a cargo has multiple motors, it is more likely to bind nearby filaments, move processively along a filament due to multiple motors binding to it, and bind to multiple filaments simultaneously. When a cargo is bound to two filaments pointed in opposite directions, forces generated by the motors walking in opposite directions serve to slide the filaments relative to one another. This mechanism is most relevant when the filaments are well mixed at short times, and it serves to locally separate filaments by polarity. Fluctuations that lead to multiple cargoes being near one another further promote the local sorting of filaments, and lead to regions of low filament density with locally ordered filaments nearby, where the plus ends are oriented toward the cargoes. These regions serve as local “traps” for additional cargoes because cargoes that bind to filaments near the cluster are directed toward the cargo-rich region.

Once a cluster of cargoes generates locally sorted filaments, if a cargo diffuses away from the cluster, it is likely to bind a filament that will transport it back. Additionally, when a filament diffuses into the cargo-rich region, motor-generated forces tend to push it back to its previous region. Both of these mechanisms stabilize local traps against perturbations as long as the filaments are sorted by polarity. The spatial distribution of cargoes, as we characterized by the radius of gyration, reveals the formation of metastable states with multiple cargo clusters at short times. The clusters are dynamic, and at longer times, the system typically coalesces into a single cargo cluster separating two filament domains. Shorter filaments suppress cargo aggregation and filament sorting, requiring more cargoes and motors to generate marked domain formation. This is likely due to less initial alignment of filaments, smaller “traps” generated by cargo-induced ordering of filaments, and increased rotational freedom of shorter filaments. Coalescence of cargo clusters is also less likely because they need to be in closer proximity to enable mixing of locally aligned filaments.

We were able to characterize polarity sorting using the average polarization in the *x*-direction because we chose an asymmetric simulation domain in which the filaments were biased by confinement to partially align with the long dimension of the system. This biased the sorting of filaments along this dimension. Our choice of domain shape was motivated by plant cells such as root epidermal cells, which commonly have a large aspect ratio. Further, the asymmetric shape provided clear signatures of sorting, which helped to reveal the underlying physics. Although we used parameters motivated by plant cells, the feedback between cargo motion and cytoskeletal organization is general and highlights mechanisms that apply in other contexts as well. Some of the results presented here are reminiscent of experimental observations. The emergence of filament-rich regions with the plus ends of filaments pointing toward a cargo cluster is similar to asters, which consist of radially oriented filaments with their ends tightly clustered together around a motor-rich region (20, 57). Aster formation has been attributed to motors that dwell at filament ends, but in this work, we show that motors that do not dwell at filament ends lead to the formation of polarity-sorted domains in confined systems. We also observe that cargoes preferentially cluster near the center of the simulation domain when the number of motors is sufficiently large. This phenomenon is similar to the preferential aggregation of myosin-II motors at the midplane of elongated actin droplets, which are liquid droplets composed of densely packed short actin filaments (58).

We focused on a system consisting solely of filaments and cargo-associated molecular motors in order to better understand their mutual impact on one another. This provides a reference point from which to better understand the addition of other important biological features, such as crosslinking proteins, anchoring of actin to the plasma membrane, and varied system shapes and sizes. These features, which are relevant in real cells, are likely to generate more complex behavior and will be interesting to explore in future studies. It will also be interesting to make new connections between simulations and experiments, for example by modulating the recruitment of motors to organelles in cells and by developing new reconstituted systems in which both cargoes and filaments are mobile.

## AUTHOR CONTRIBUTIONS

O.H.A. and S.M.A. designed research. O.H.A. performed research. O.H.A. and S.M.A. analyzed data and wrote the manuscript.

## ACKNOWLEDGMENTS

This work was supported by the National Science Foundation (MCB-1715794 and MCB-2334517).

