## Supplemental information for "Polarity sorting of actin filaments by motor-driven cargo transport"

| Symbol | Description (units) | Value |
|---|---|---|
| | **Actin filaments** | |
| $N_f$ | number of filaments | 100, 200 |
| $L$ | total filament length ($\mu$m) | 1000 |
| $l$ | Number of links per filament $= L/N_f$ | 10, 5 |
| $l_a$ | link rest length ($\mu$m) | 1 |
| $k_a$ | stretching force constant (pN/$\mu$m) | 1 |
| $\kappa_B$ | bending modulus (pN·$\mu$m$^2$) | 0.068 |
| | **Cargo and motors** | |
| $N_c$ | number of cargoes | 10, 20, 40, 80 |
| $N_m$ | number of motors per cargo | 1, 2, 4, 8 |
| $r_c$ | cargo radius ($\mu$m) | 0.25 |
| $l_m$ | motor rest length ($\mu$m) | 0.35 |
| $k_m$ | motor stiffness (pN/$\mu$m) | 10 |
| $k_m^{\text{on}}$ | motor attachment rate (s$^{-1}$) | 5 |
| $k_{0,m}^{\text{off}}$ | unloaded motor detachment rate (s$^{-1}$) | 5 |
| $k_m^{\text{end}}$ | barbed end detachment rate (s$^{-1}$) | 100 |
| $F_d$ | characteristic detachment force (pN) | 1 |
| $v_0$ | unloaded motor speed ($\mu$m/s) | 7 |
| $F_s$ | stall force of myosin (pN) | 0.5 |
| | **Environment and simulation details** | |
| $\Delta t$ | simulation timestep (s) | 0.00002 |
| $t_F$ | simulated time (s) | 500 |
| $T$ | temperature (K) | 298 |
| $\nu$ | dynamic viscosity (Pa·s) | 0.001 |

Table 1: Table of parameters used in simulation.

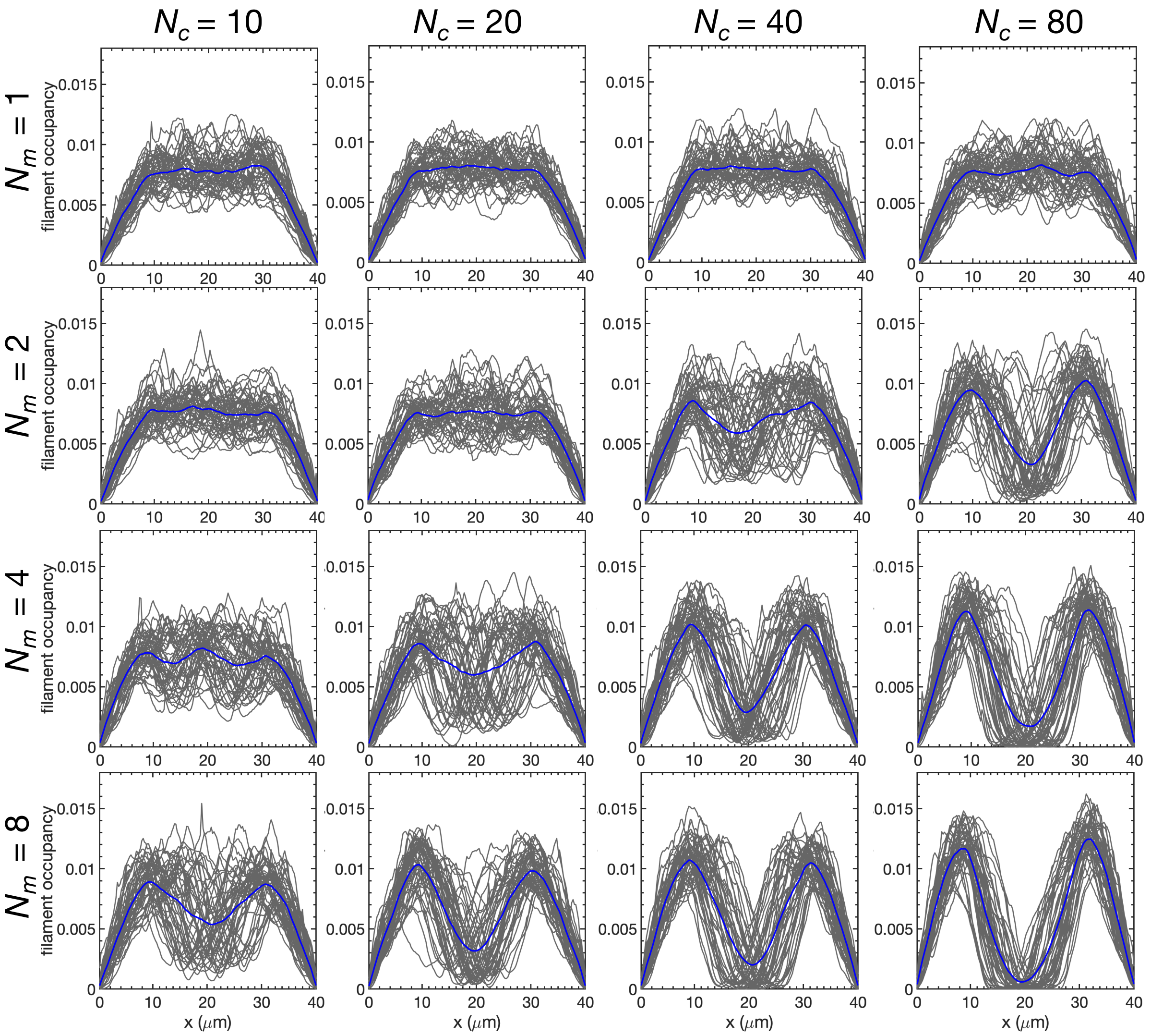


Figure S1: Filament occupancy as a function of position along the long dimension of the simulation box ($x$). Results are shown for 10 $\mu$m filaments and different values of $N_c$ and $N_m$. Each grey curve is the average occupancy over the last 50 s of an independent trajectory. The mean across all trajectories is shown in blue.

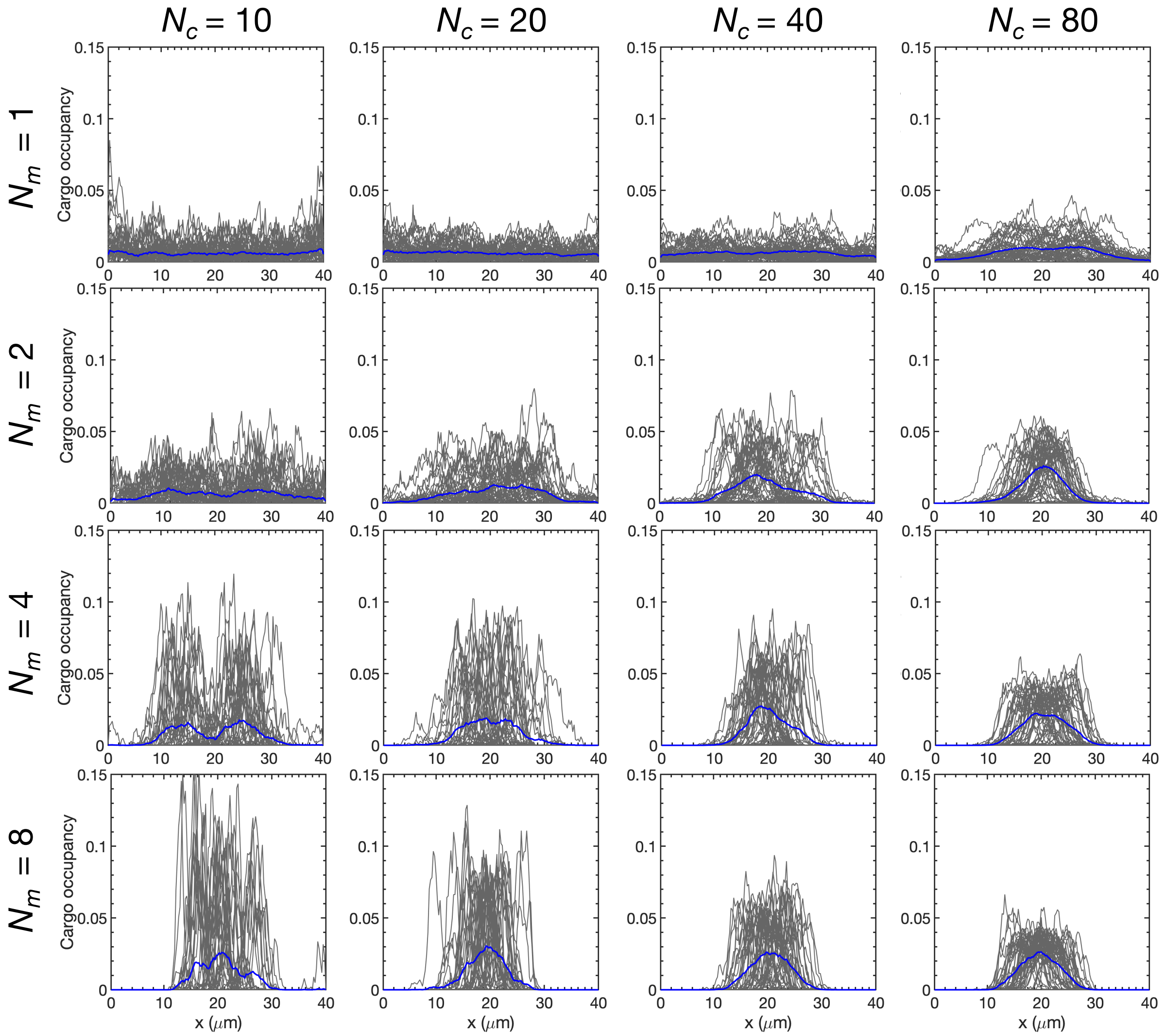


Figure S2: Cargo occupancy as a function of position along the long dimension of the simulation box ($x$). Results are shown for 10 $\mu$m filaments and different values of $N_c$ and $N_m$. Each grey curve is the average occupancy over the last 50 s of an independent trajectory. The mean across all trajectories is shown in blue.

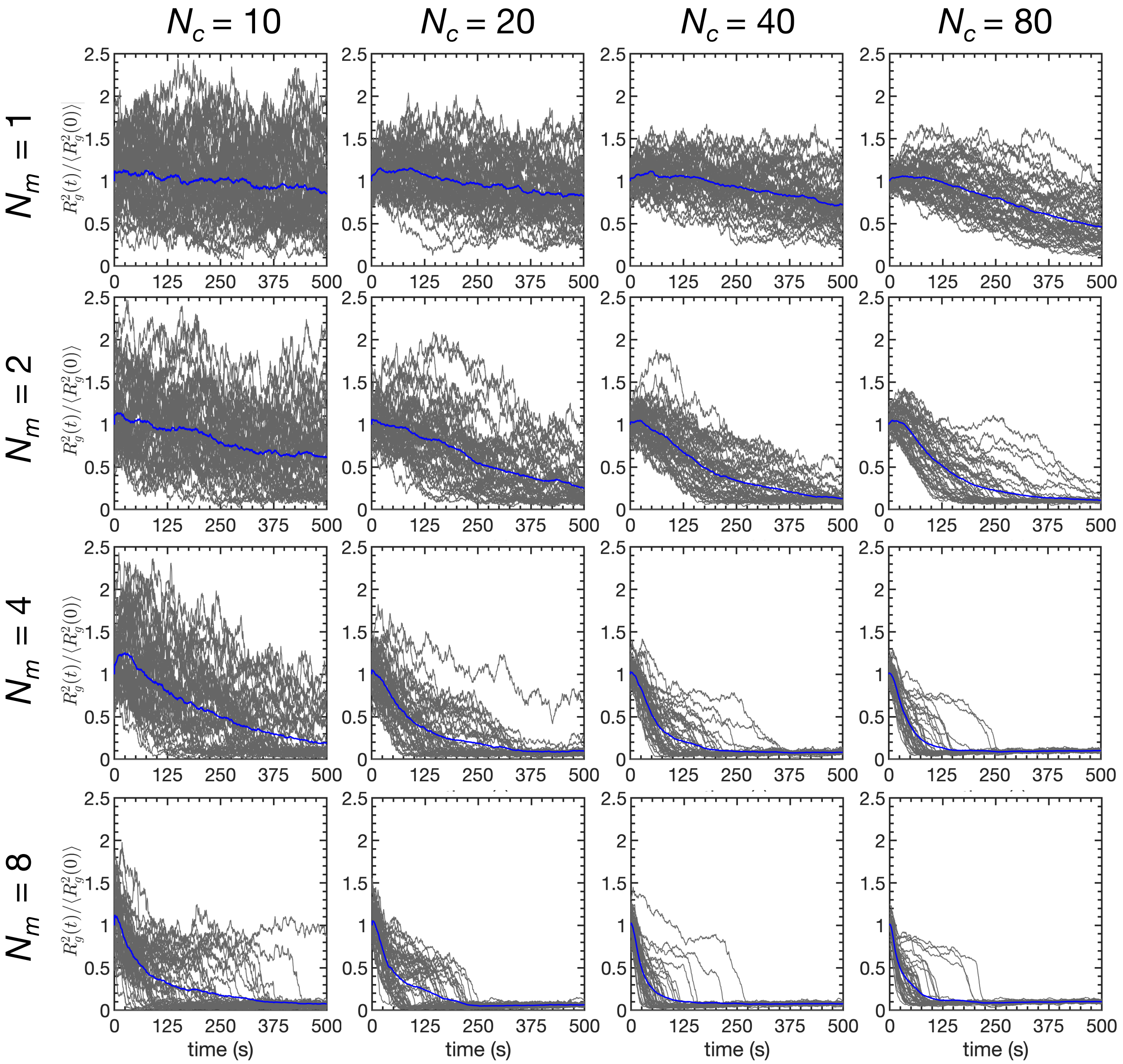


Figure S3: Normalized square radius of gyration for 10 $\mu$m filaments and different values of $N_c$ and $N_m$. Each individual trajectory is shown in grey. The mean across all trajectories is shown in blue.

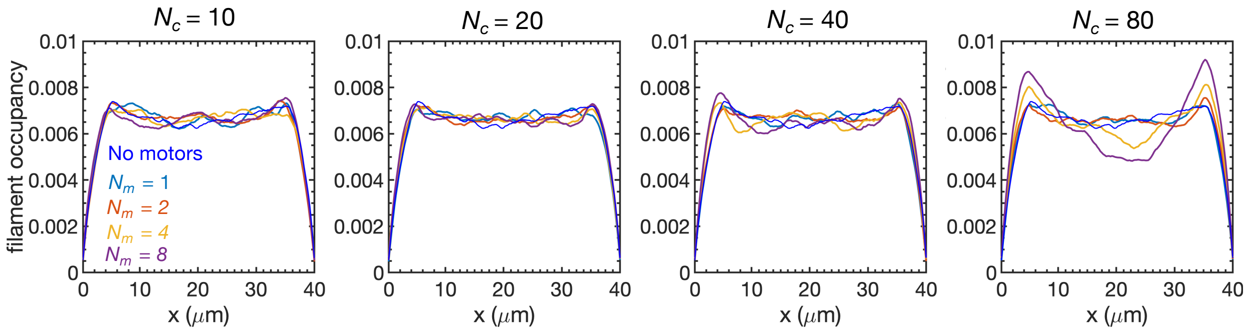


Figure S4: Mean filament occupancy as a function of position along the long dimension of the simulation domain ($x$) for networks with 5 $\mu$m filaments and different values of $N_c$ and $N_m$.

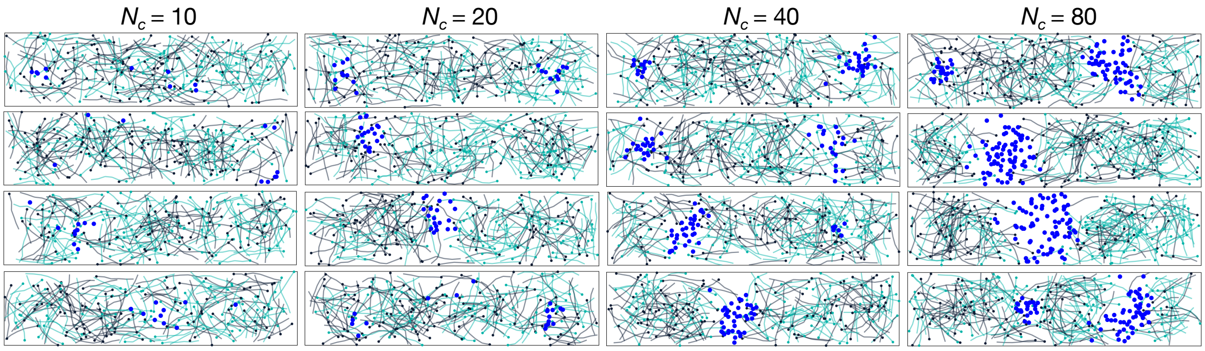


Figure S5: Final snapshots of independent trajectories with 5 $\mu$m filaments and $N_m = 8$. Four representative replicates are shown for each value of $N_c$. Filaments are colored according to their initial orientation, with left-facing filaments shown in green and right-facing filaments shown in charcoal. Cargoes are depicted by blue circles. Individual motors are not shown.

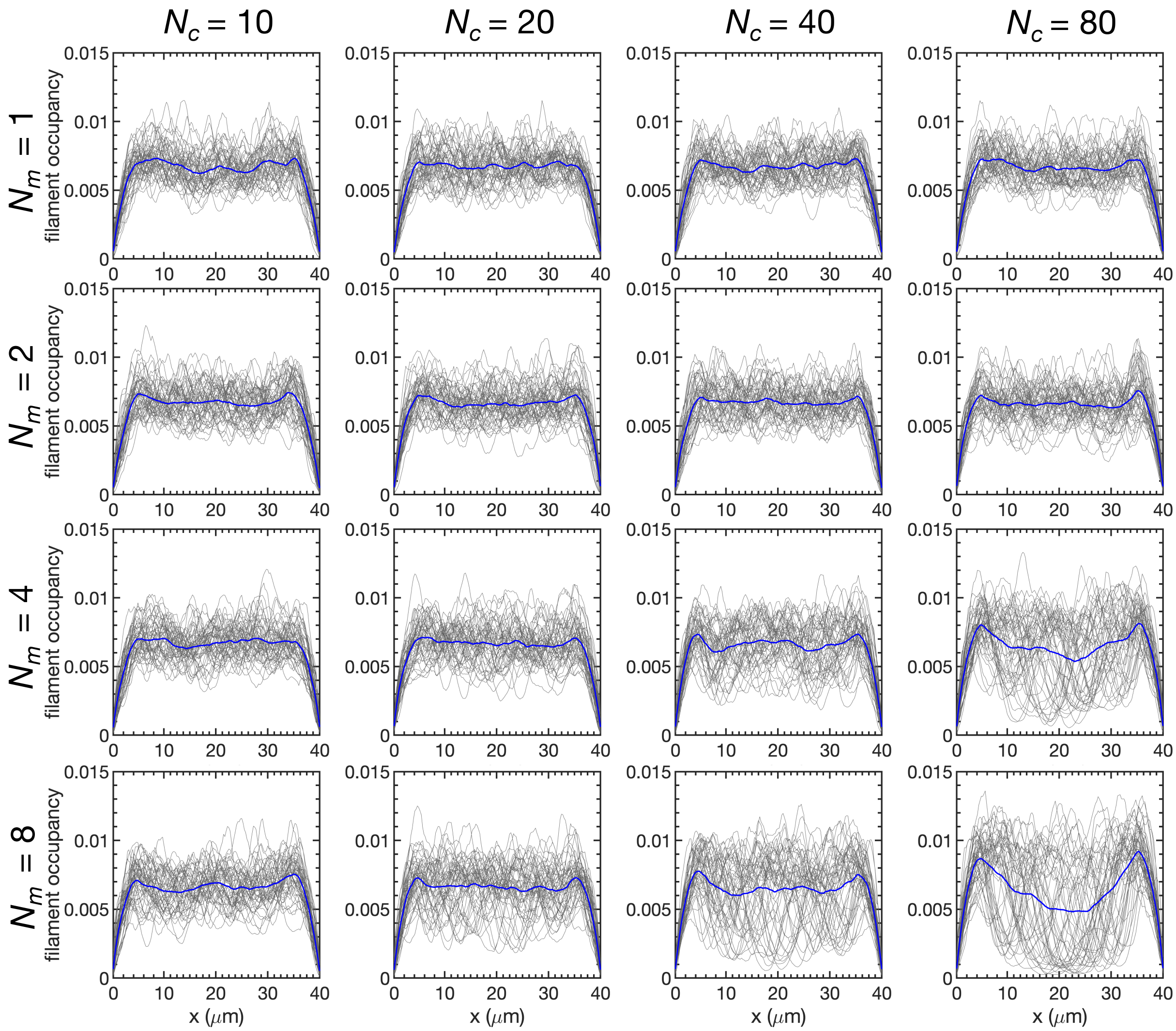


Figure S6: Filament occupancy as a function of position along the long dimension of the simulation box ($x$). Results are shown for 5 $\mu$m filaments and different values of $N_c$ and $N_m$. Each grey curve is the average occupancy over the last 50 s of an independent trajectory. The mean across all trajectories is shown in blue.

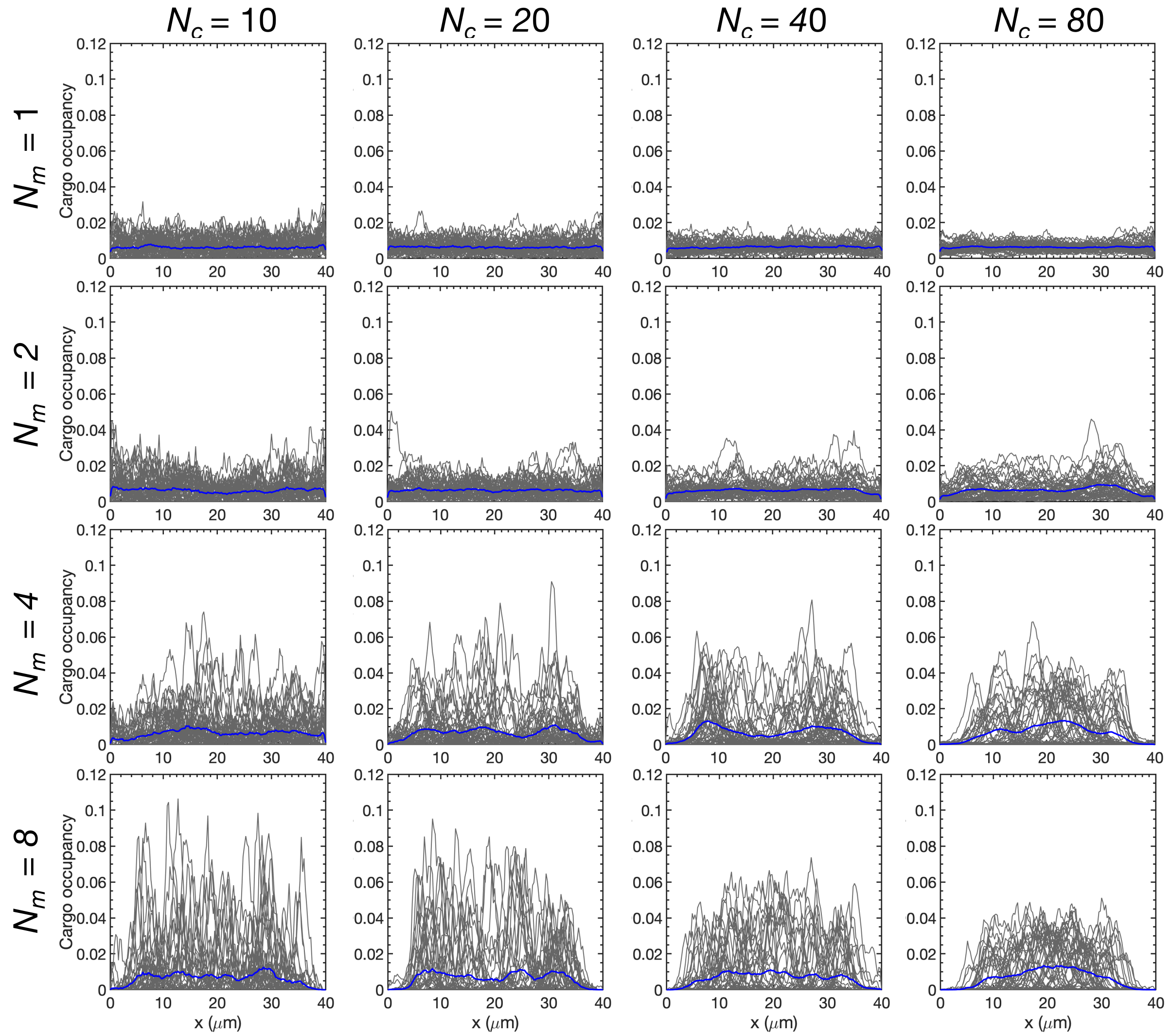


Figure S7: Cargo occupancy as a function of position along the long dimension of the simulation box ($x$). Results are shown for 5 $\mu$m filaments and different values of $N_c$ and $N_m$. Each grey curve is the average occupancy over the last 50 s of an independent trajectory. The mean across all trajectories is shown in blue.

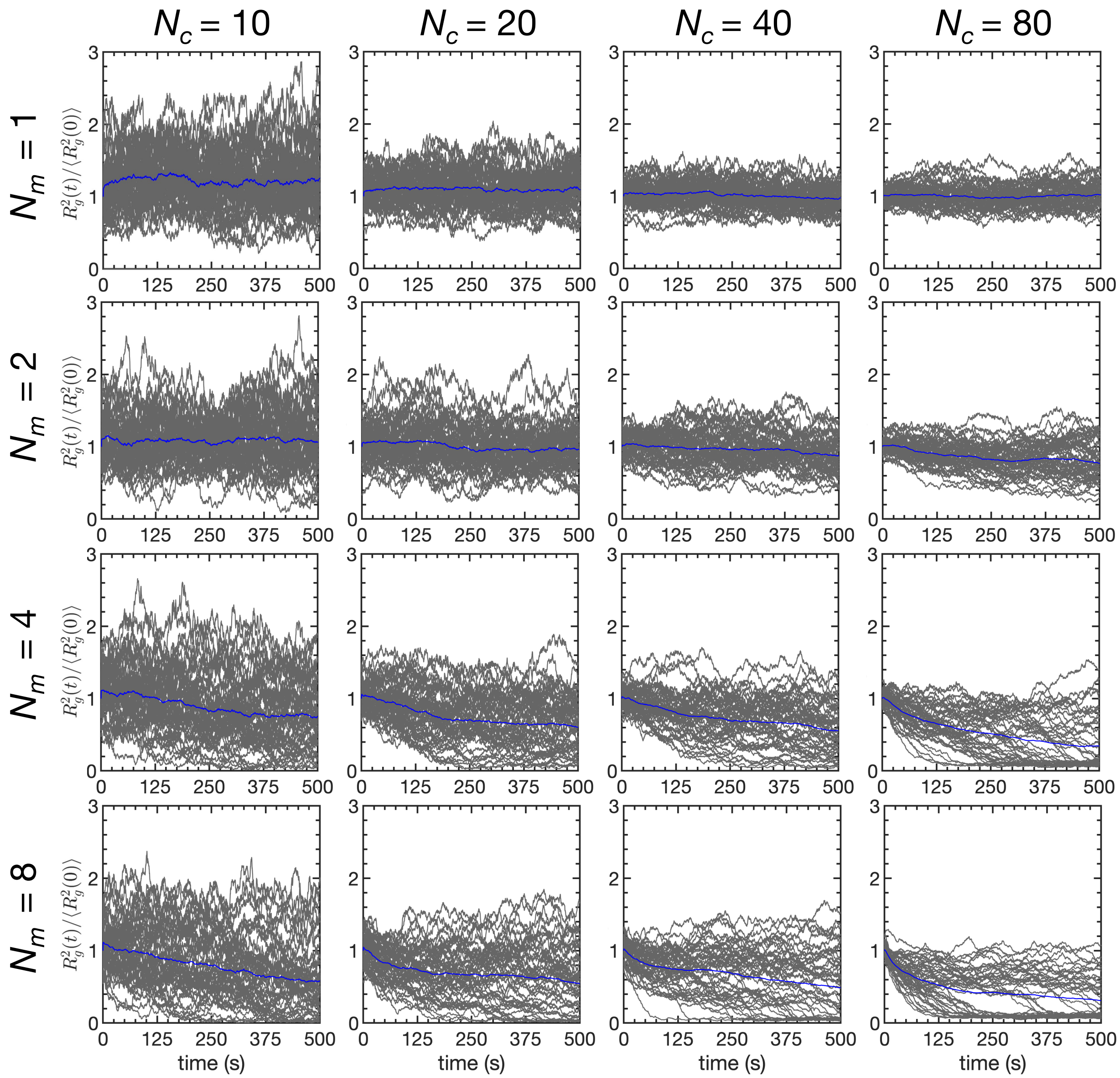


Figure S8: Normalized square radius of gyration for 5 $\mu$m filaments and different values of $N_c$ and $N_m$. Each individual trajectory is shown in grey. The mean across all trajectories is shown in blue.
